# Early-session facial dynamics predict executive-function response to digital cognitive interventions in older adults: development and external validation

**DOI:** 10.1101/2025.07.29.667519

**Authors:** Yang Merik Liu, Adam Turnbull, Meishan Ai, Kathi L. Heffner, Cristiano Tapparello, Yu Zhang, Ehsan Adeli, Feng Vankee-Lin

## Abstract

Digital cognitive interventions can improve cognition in older adults, but individual differences in transfer to executive function (EF) remain substantial and are difficult to anticipate before treatment completion. We developed a compact Facial Signature Score (FSS) from passive webcam recordings collected during therapeutic cognitive challenges in BREATHE (NCT04522791; development, N=87) and evaluated it in PSOPT (NCT06005038; external validation, N=39), an independent trial using a different intervention format. The FSS discriminated EF responders under nested cross-validation in BREATHE (AUROC=0.77) and generalized to PSOPT without feature reselection or reweighting (AUROC=0.78). The FSS was associated with continuous EF change in BREATHE (Spearman ρ=0.49, p<0.001) and after outcome-independent recalibration in PSOPT (ρ=0.32, p=0.045). Predictive performance was retained when the signature was computed from the first 25% and 50% of intervention sessions, and the FSS added value beyond baseline characteristics and conventional behavioral measures. Passive facial dynamics during therapeutic cognitive challenge may serve as a scalable intervention-process marker for early assessment of cognitive response among older adults.

## Introduction

Cognitive interventions provide a scalable approach to preserving or improving cognitive function in older adults, including individuals with mild cognitive impairment (MCI).^1^ Among these interventions, computerized cognitive training (CCT) targeting processing speed and attention (PS/A) has shown consistent benefits for the trained cognitive domains, with durable effects observed following booster sessions.^2^ Digital cognitive interventions have also increasingly incorporated additional components to CCT, such as resonance-frequency breathing or guided imagery relaxation, as well as adaptive closed-loop training paradigms intended to personalize intervention delivery.^3–5^ Despite differences in implementation, these approaches share the use of repeated cognitive challenge to promote cognitive improvement.^6^ Although PS/A-based CCTs can improve trained abilities, transfer to executive function (EF), a key determinant of adaptative functioning and independence in everyday life, remains heterogeneous across individuals.^7^ This variability suggests that transfer depends not only on exposure to cognitive training but also on individual differences in how participants engage with and adapt to repeated cognitive challenges during intervention.^8,9^ An important challenge, therefore, is to identify which individuals are likely to show EF improvement and whether this response can be detected early enough to support personalized intervention delivery.

Existing response-prediction approaches have primarily relied on pretreatment demographic, cognitive, neuroimaging, or functional characteristics.^10,11^ For example, a recent machine-learning analysis of the ACTIVE trial used baseline participant characteristics to predict responsiveness to cognitive training but achieved only modest predictive performance, particularly for cognitive outcomes more distal from the trained tasks.^11,12^ Baseline measures characterize an individual’s predisposition before treatment begins but provide little information about how that individual responds during repeated therapeutic cognitive challenges. This distinction may be particularly relevant to predicting transfer to EF. Such transfer is likely influenced by dynamic intervention-processes characteristics, such as motivation, sustained attention, cognitive flexibility, and self-regulation, that unfold across repeated cognitive challenges.^10,13^ These characteristics are not directly reflected by conventional training measures, such as adherence, reaction time, accuracy, and training dose,^13^ which primarily quantify exposure to, or performance on, the trained tasks rather than how individuals engage with the therapeutic process itself.

Passive behavioral sensing offers a scalable way of capturing intervention-process information without interrupting treatment or increasing participant burden.^14,15^ Consumer-grade webcams enable continuous recording of facial behavior while participants engage in standardized cognitive challenges that share core cognitive demands with the interventions themselves.^16^ Facial action units (AUs), which encode anatomically based facial muscle movements, provide an interpretable representation that can be quantified to characterize behavioral dynamics across repeated cognitive challenges.^17,18^ Facial dynamics expressed during these challenges may provide observable correlates of engagement-related states with the therapeutic process, such as fatigue and attentional variation, and could potentially reflect broader self-regulatory or adaptive responses relevant to EF transfer.^13,19^ Because these signals are acquired while individuals actively engage with therapeutic cognitive challenges, they have the potential to capture intervention-process information that is unavailable from baseline characteristics alone.^20^ Previous studies have shown that facial video contains information related to cognitive and behavioral state, including fatigue during cognitive-training sessions and differences among individuals with MCI, dementia, and cognitively unimpaired older adults.^16,21,22^ However, this work has focused primarily on estimating concurrent state or classifying existing cognitive status. Whether facial dynamics recorded during therapeutic cognitive challenges contain information that predicts subsequent EF improvement remains unknown. It is also unclear whether any predictive information generalizes across different cognitive intervention formats or instead reflects characteristics of a specific training program.

In this study, we analyzed facial dynamics collected during two independent randomized controlled trials (RCTs) of older adults with or at risk for cognitive impairment. The trials used different cognitive-intervention formats centered on PS/A-based CCT. Using automated computer vision, we extracted multiscale temporal features from facial AUs recorded during intervention sessions and developed a compact facial response signature in the BREATHE trial.^3,17^ We then externally validated the signature in the independent PSOPT trial without feature reselection or outcome-informed model updating. We addressed four objectives: (1) to determine whether passive facial dynamics recorded during therapeutic cognitive challenges predicted EF improvement and generalized across cohorts using different intervention formats; (2) to evaluate whether a facial signature developed from the full intervention course retained predictive performance when computed using only early-session data; (3) to determine whether the facial signature provided incremental predictive value beyond baseline characteristics and conventional behavioral measures; and (4) to assess the robustness of the facial signature across training tasks and randomized intervention arms.

## Methods

### Study cohorts and participants

We analyzed two independent randomized controlled trial (RCT) cohorts of older adults participating in digital cognitive interventions (**Figure 1A**). The development cohort (BREATHE; ClinicalTrials.gov identifier: NCT04522791; registered August 21, 2020) included cognitively unimpaired older adults and older adults with mild cognitive impairment (MCI).

**Figure 1.**
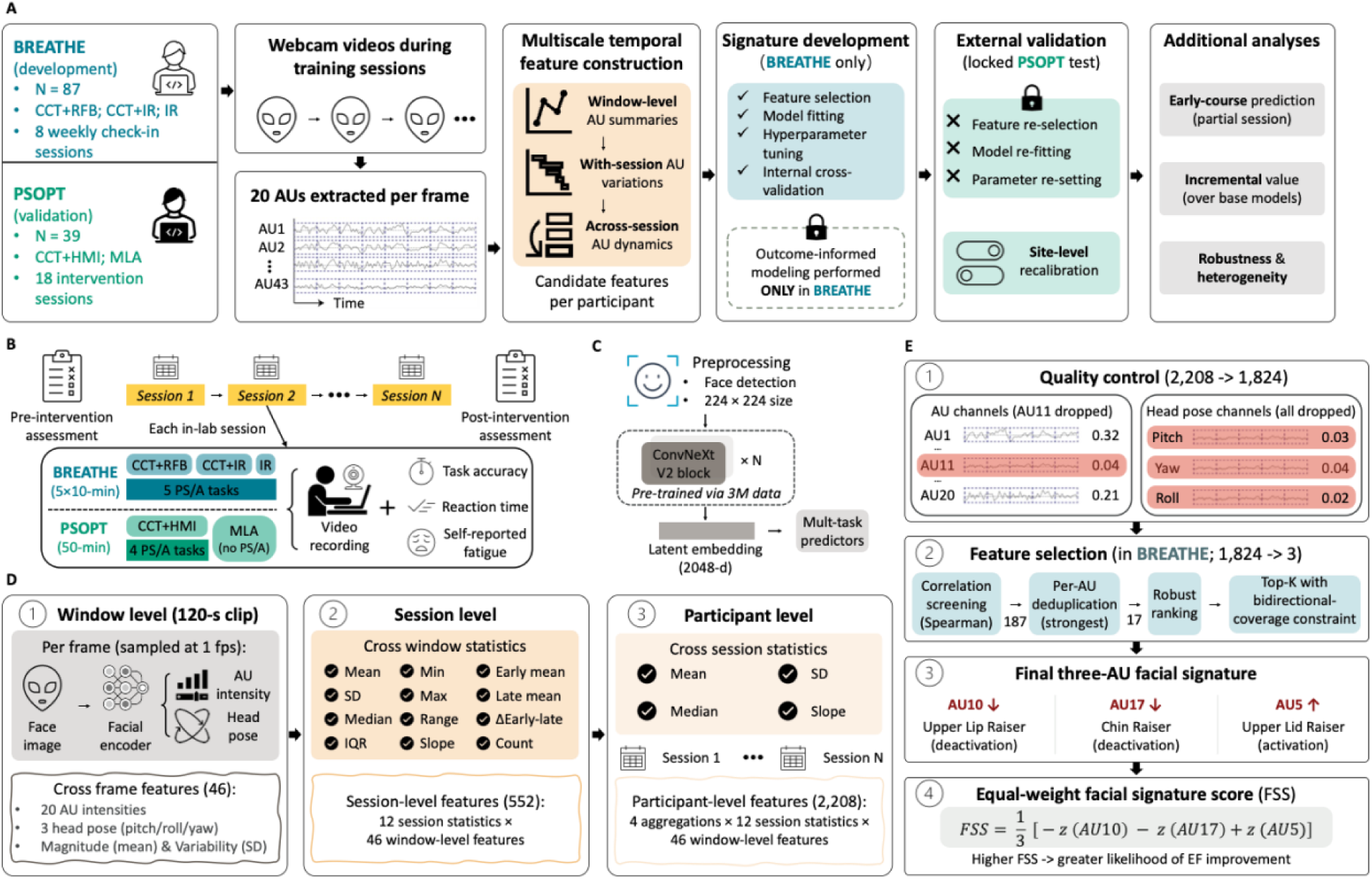
(A) Overall pipeline for model development and external validation. All outcome-informed steps, including facial-feature extraction, feature selection, direction alignment, feature standardization, and FSS construction, were performed in BREATHE; PSOPT was used only for external validation and outcome-independent recalibration. (B) Study protocol and video-recording structure across repeated intervention sessions. (C) Facial encoder used to extract frame-level AU intensities and head-pose estimates from training-session video. (D) Multiscale construction of AU-based features across window-level, within-session, and across-session timescales. (E) Feature-selection cascade leading to the final facial signature, defined from sign-aligned AU10, AU17, and AU5 features.

Participants with MCI were randomized to one of three intervention conditions: processing speed/attention (PS/A)-based computerized cognitive training combined with resonance-frequency breathing (CCT+RFB), PS/A-based CCT combined with imagery-guided relaxation (CCT+IR), or imagery-guided relaxation alone (IR). Cognitively unimpaired participants received CCT+RFB. During each of up to eight weekly in-laboratory sessions, participants completed standardized 50-minute PS/A cognitive challenges, including multiple object tracking (MOT), rapid serial visual presentation (RSVP), visual search (Search), visual sweeps (Sweeps), and useful field of view (UFOV), while facial video was continuously recorded (see **Supplementary Methods 1** and **Table S1A**). These cognitive challenges shared core cognitive demands with, but were not identical to, the intervention tasks.

The external-validation cohort (PSOPT; ClinicalTrials.gov identifier: NCT06005038; registered August 18, 2023) included older adults with at least one dementia risk factor who were randomized to PS/A-based CCT integrated with a closed-loop human–machine interface (CCT+HMI) or a mental leisure activities control (MLA). The PS/A cognitive challenges included motion of target, sound sweeps, mixed signal, and delayed task switching (**Table S1A**). Facial video was recorded throughout all intervention sessions (up to 18 sessions).

Both trials were conducted in accordance with the principles of the Declaration of Helsinki. The BREATHE trial was approved by the University of Rochester Research Subjects Review Board, serving as the single Institutional Review Board (sIRB), with Stanford University participating as a study site (ethics approval no. 00004727). The PSOPT trial was approved by the Stanford University Institutional Review Board (ethics approval no. 71235). All participants provided written informed consent before study participation. Primary analyses included participants with usable facial recordings and complete executive-function assessments (BREATHE, N=87; PSOPT, N=39; **Supplementary Table S2** and **Figure S1** for participant flow and recording quality). Participants’ characteristics are summarized in **Table 1**.

**Table 1.** Participant characteristics of the development and external cohorts.

| Characteristic | BREATHE (development) | PSOPT (external) |
| --- | --- | --- |
| N | 87 | 39 |
| Age, years — mean ± SD | 72.5 ± 7.2 | 76.5 ± 7.8 |
| Female — n (%) | 55 (63%) | 22 (56%) |
| Education, years — mean ± SD | 15.9 ± 2.5 | 16.8 ± 2.5 |
| MoCA — mean ± SD | 24.8 ± 2.5 | 25.2 ± 3.0 |
| Executive-function responders — n (%) | 54 (62%) | 27 (69%) |
| Data collection sessions — median (IQR) | 8 (7–8) | 18 (17–18) |
| Randomized allocation — n | CCT+RFB: 51<sup>a</sup>; CCT+IR: 16; IR: 20 | CCT+HMI: 20; MLA: 19 |

<sup>a</sup> PS/A+RFB includes all 28 cognitively unimpaired participants and 23 with MCI; other two arms are MCI only.
<sup>b</sup> MoCA, Montreal cognitive assessment; RFB, resonance-frequency breathing; IR, imagery-guided relaxation; MLA, mental leisure activities.

### Measures

The primary cognitive outcome was an EF composite score, computed at baseline and after intervention as the mean of within-cohort z-standardized EF task scores from NIH EXAMINER battery.^23^ The primary outcome was binary EF response, defined as any measurable EF improvement (EF change > 0). Continuous EF change was analyzed as a secondary outcome, with baseline EF included as a covariate in adjusted analyses.

Conventional behavioral measures were collected during the same cognitive challenge sessions as facial video recording (**Figure 1B**). These included self-reported fatigue (Visual Analogue Scale, pre- and post-session), task accuracy, and reaction time. Reaction-time measures were standardized within each training task, averaged across tasks within session, and averaged across sessions. For incremental analyses, these measures were summarized as mean within-session fatigue change, mean task accuracy, and intra-individual reaction-time variability (IIV-RT; the within-person coefficient of variation of reaction time).

Baseline characteristics consisted of age, sex, years of education, and an Alzheimer’s disease-signature cortical thickness (ADSCT) index. ADSCT was derived from baseline structural MRI as a neuroanatomical summary of cortical thickness in AD-vulnerable regions, with lower values indicating greater neurodegeneration.^24^ The baseline EF composite was additionally used as a covariate in a sensitivity model.

See summaries of measurement definitions in **Supplementary Table S1B**.

### Prediction model development and validation

#### Overview

All predictive model development was performed exclusively in the BREATHE cohort to maintain complete separation between model development and external validation. This included facial-feature extraction, feature selection, direction alignment, feature standardization, and FSS construction. The independent PSOPT cohort was reserved exclusively for external validation and was not used for any outcome-informed analytical decisions. An outcome-independent sensitivity analysis recalibrated feature standardization using the PSOPT predictor distribution while keeping the selected features, direction alignment, and equal-weight scoring rule unchanged. **Figure 1A** illustrates the overall analytical workflow.

#### Facial feature extraction

Facial action units (AUs) were extracted frame by frame from training-session videos using py-feat (version 2.0.2; **Figure 1C**), yielding intensity estimates for 20 facial muscles.^25^ Frames without reliable face detection were excluded. To characterize facial dynamics throughout intervention, AU intensities were summarized hierarchically across three temporal scales: analysis windows (120 s), intervention sessions, and participants. Summary measures captured the magnitude, variability, and temporal organization of facial behavior within and across sessions (**Figure 1D**). This multiscale representation generated 2,208 participant-level candidate features (**Supplementary Methods 2, Table S3**). AU channels exhibiting near-constant output, consistent with detector saturation, were removed through a predefined quality-control procedure in the development cohort.

#### Development of the facial signature

Feature selection was performed only in BREATHE using continuous EF change as the development outcome, and the full selection procedure was repeated within each outer cross-validation fold to prevent information leakage. Candidate features were first screened by their Spearman rank correlation with EF change, retaining only features that reached p < 0.05. A rank correlation was used because the EF-change outcome was strongly right-skewed (Shapiro–Wilk p < 10⁻⁶) and most candidate features were non-normal.

This screen served only as a feature-filtering step, and development-stage p values were treated as screening statistics rather than confirmatory inference. Among features from the same AU, the strongest feature was retained by absolute biweight midcorrelation. The remaining AU-level features were ranked by the same robust statistic, and the final three-feature signature was selected using a predefined directional-coverage rule requiring both positive and negative associations (**Figure 1E**). Signature size (K=3) was fixed *a priori* using an events-per-variable criterion of at least 10 outcome events per predictor. The equal-weight Facial Signature Score (FSS) was computed as the mean of the three z-standardized, sign-aligned features, with higher values indicating greater predicted likelihood of EF improvement. The FSS was evaluated as the primary predictor for EF-responder discrimination and association with continuous EF change.

#### FSS development and internal validation

Binary EF response was predicted directly from the FSS. Because the FSS combines the three sign-aligned standardized features using fixed equal weights, no feature-specific prediction weights were fitted, and discrimination was quantified as the AUROC of the FSS itself. To place predictions on a probability scale for calibration, a univariable logistic link between EF response and the FSS was fitted in BREATHE and then locked {logit(*p*) = 0.61 + 1.60 × FSS}. Because this link is strictly monotonic, AUROC is identical on the FSS and probability scales. A prespecified classification threshold of 0.50 predicted probability was used for binary classification, and no hyperparameter search was performed. Internal model performance was estimated using fully nested leave-one-subject-out (LOSO) cross-validation. Within each outer fold, all outcome-informed procedures, including feature screening, within-AU deduplication, robust ranking, direction alignment and feature standardization, were repeated using only the training participants to prevent information leakage. Feature-selection stability was evaluated by recording selection frequencies across outer folds. To quantify optimism arising from feature search, the complete development pipeline was repeated across 1,000 participant-level outcome permutations. Additional sensitivity analyses are described in the **Supplementary Methods 3-4** and the locked model specification is given **in** Supplementary Table S5A.

#### External validation

External validation was performed by applying the locked FSS to the independent PSOPT cohort without feature reselection or signature reconstruction. Discrimination, calibration and the association with continuous EF change were all evaluated from the frozen FSS, with calibration computed from the locked probability link. External discrimination was additionally tested by a conditional permutation test (1,000 permutations of PSOPT outcome labels with the signature and link held fixed). Because the cohorts differed in participant characteristics, intervention delivery, and recording conditions, an outcome-independent sensitivity analysis recalibrated only feature-standardization parameters using the PSOPT predictor distribution while leaving feature definitions, direction alignment, the equal-weight scoring rule, the locked probability link and the classification threshold unchanged.

#### Prediction across intervention-course intervals

To evaluate how predictive performance changed as progressively more intervention data became available, the facial signature derived from the full intervention course was reconstructed using first 25% and first 50% of each participant’s intervention course. The three selected facial features and their direction alignment were held fixed across all intervention-course intervals. For each interval, feature-standardization parameters were estimated and the same equal-weight scoring rule was applied, with nothing refitted. Because the interval analysis recomputes the fixed feature definitions from window-level data, it constitutes a distinct feature-construction pipeline from the primary analysis. Full-course interval estimates are therefore close to, but not identical with, the primary estimates.

### Sensitivity analyses

#### Incremental value

Incremental analyses tested whether the FSS added information about EF change beyond interpretable base models in BREATHE. The behavioral model comprised fatigue change, task accuracy, and IIV-RT. The baseline-characteristics model comprised age, sex, education and ADSCT, with baseline EF added in a third sensitivity model. Incremental continuous value was quantified by change in R² and a one-degree-of-freedom partial F test. Classification value was evaluated using fixed-signature LOSO AUROC and bootstrap ΔAUROC confidence intervals. Note, ΔR² is an in-sample statistic for a continuous outcome and ΔAUROC a cross-validated statistic for a binary outcome. These are distinct estimands and are not expected to agree in significance. The incremental analysis was repeated across early-course intervals, and PSOPT analyses adjusted for age, sex, education, and baseline EF only because ADSCT and conventional behavioral measures were unavailable externally.

#### Intervention-arm heterogeneity

In each randomized cohort, we tested whether the FSS association with EF improvement depended on training arm. In BREATHE, post-intervention EF was modeled as a function of baseline EF, FSS, intervention arm, and the FSS-by-arm interaction. The omnibus interaction was the treatment-heterogeneity test, and the FSS main effect was the prognostic estimate. The full model included all BREATHE participants (both cognitively unimpaired and MCI; N=87). Because only the CCT+RFB arm enrolled cognitively unimpaired participants, the FSS-by-arm interaction was additionally repeated restricted to participants with MCI (N=59, across all three arms). An FSS-by-cognitive-status interaction was tested within the CCT+RFB arm (28 cognitively unimpaired, 23 MCI). Analogous models were estimated in PSOPT. All arm models were fit by ordinary least squares with heteroskedasticity-consistent robust standard errors to reduce sensitivity to nonconstant residual variance in these modest-sized subgroup analyses.

#### Robustness across cognitive challenges

In BREATHE, we tested whether the facial signature depended on any individual cognitive challenge. For leave-one-task-out analyses, we recomputed the three fixed feature definitions after removing all video segments from one task type at a time, using the remaining tasks to construct the FSS. For single-task analyses, we recomputed the same three feature definitions using video segments from one task type only. Reconstructions were evaluated by the AUROC of the reconstructed FSS and its Spearman correlation with EF change. Equivalent task-specific analyses were not possible in PSOPT because recordings were not separately task-labeled.

### Statistical analysis

Classification performance was summarized using AUROC. Calibration was summarized using Brier score, calibration slope, and calibration intercept. Continuous FSS–EF associations were summarized primarily using Spearman rank correlations, reflecting the use of the FSS as a response-ranking marker. Participant-level bootstrap resampling with 2,000 replicates was used to estimate 95% confidence intervals unless otherwise specified. For internal cross-validation metrics, participants and their out-of-fold predictions were resampled together. For paired model comparisons, participants were resampled with observed labels and predictions from both models. External confidence intervals were estimated by resampling PSOPT participants.

Empirical one-sided permutation p values were calculated as *p* = (*b* + 1)/(*B* + 1), where b was the number of permuted statistics at least as extreme as the observed statistic and B the number of permutations. These tests were one-sided because the alternative hypothesis was directional, namely discrimination above chance. Other tests were two-sided with a nominal significance level of p < 0.05.

Analyses used Python 3.11.15 with scikit-learn 1.7.2, statsmodels 0.14.6, SciPy 1.11.4, NumPy 1.26.4, pandas 2.3.3, and py-feat 2.0.2, with random seed 42. Detailed parameter settings, analysis provenance, sample flow, and sensitivity and recording-quality analyses are provided in the **Supplementary Methods 2–4**, **Table S2** and **Figure S1**.

## Results

### Objective 1. Development and validation of an intervention-process facial signature

#### Development of the facial signature in BREATHE

We first developed a facial signature associated with subsequent EF improvement using the BREATHE development cohort (N=87). Automated facial coding generated 2,208 participant-level summaries of facial AU dynamics across intervention sessions. Following quality-control procedures and outcome-informed feature selection (**Methods**), three features representing three facial muscle movements were retained (**Figure 2A**; **Supplementary Table S4A–B**). Two lower-face features captured between-session instability in within-session dynamics: the across-session variability of early-to-late change in AU10 (Upper Lip Raiser) and the across-session variability of within-session fluctuations in AU17 (Chin Raiser). Higher values of both features were associated with less subsequent EF improvement. The third feature represented the across-session median of within-session median in AU5 (Upper Lid Raiser), with greater values associated with greater EF improvement. These three features were consistently reselected across nested cross-validation folds (selection frequency 97%–100%; **Supplementary Table S4C**). The FSS discriminated participants with versus without EF improvement under fully nested LOSO cross-validation, in which feature selection, direction alignment and standardization were re-derived within each fold (AUROC=0.77, 95% CI: 0.66–0.86; **Figure 2B**). A full-pipeline permutation analysis that repeated the entire selection and scoring procedure across 1,000 outcome permutations confirmed above-chance discrimination (p=0.030; **Supplementary Figure S2A**). Using the fixed full-cohort signature, higher FSS values were associated with greater continuous EF improvement (Spearman ρ=0.49, p<0.001; **Figure 2C**).

**Figure 2.**
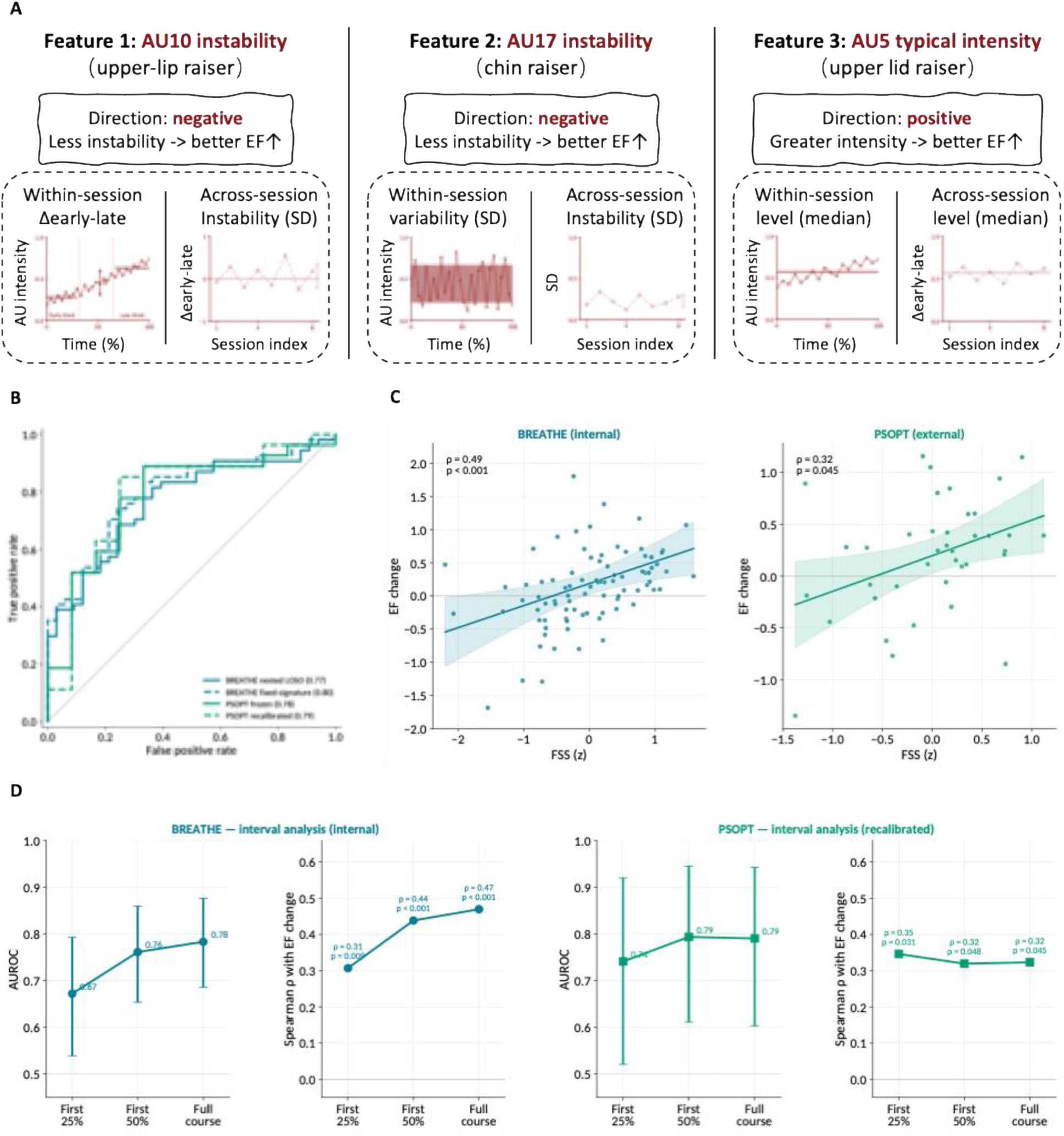
Facial-signature composition and predictive performance across development, external validation, and early-course intervals. (A) FSS comprised three AU-based features. (B) AUROC curves for EF responder prediction. BREATHE performance is shown for fully nested LOSO validation and fixed-signature evaluation; PSOPT performance is shown for the frozen FSS and outcome-independent site-level recalibration. (C) Associations between FSS and ΔEF in BREATHE and PSOPT. Lines show fitted linear trends with 95% confidence bands. (D) Early-course performance in BREATHE and PSOPT using the first 25%, first 50%, and complete intervention course.

#### External validation in PSOPT

We next evaluated whether the facial signature generalized to an independent cohort employing a different digital cognitive intervention format. The locked facial signature developed in BREATHE was applied to the independent PSOPT cohort (N=39) without feature reselection, model refitting, or threshold modification. Under this strictly locked validation, the frozen FSS discriminated participants with versus without EF improvement with an AUROC of 0.78 (95% CI: 0.60–0.93; **Figure 2B**), which exceeded chance in a conditional permutation test with all parameters held fixed (p=0.002; **Supplementary Figure S2B**). The FSS showed a positive but non-significant association with continuous EF improvement (Spearman ρ=0.30, p=0.060; **Figure 2C**). Because the development and validation cohorts differed in participant characteristics, intervention implementation, and recording conditions, we additionally evaluated an outcome-independent site-level recalibration that re-estimated only feature-standardization parameters while keeping the facial signature, direction alignment and equal-weight scoring rule unchanged. Discrimination remained similar following recalibration (AUROC=0.79, 95% CI: 0.60–0.94; **Figure 2B**), and the recalibrated FSS was associated with EF improvement (Spearman ρ=0.32, p=0.045; **Figure 2C**). Calibration also improved descriptively, with the calibration slope increasing from 0.71 to 1.19, the calibration intercept decreasing from 1.33 to 0.27, and the Brier score improving from 0.25 to 0.17 (**Supplementary Table S5B, Figure S2C–D**). Although the association with the magnitude of continuous change was weaker externally, these findings indicate that responder discrimination generalized across independent cohorts.

### Objective 2. Prediction across intervention-course intervals

To determine how predictive performance changed as progressively more intervention data became available, we reconstructed the facial signature using different portions of each participant’s intervention course while keeping the selected facial features and their direction alignment fixed. Performance at each intervention-course interval was compared with the complete intervention-course reference in both the development and external-validation cohorts. In the BREATHE development cohort, predictive performance improved progressively as additional intervention sessions were incorporated. Discrimination increased from an AUROC of 0.67 using the first 25% of sessions to 0.76 using the first 50%, approaching the full-course interval estimate of 0.78 (**Figure 2D**). Similarly, the association between FSS and continuous EF improvement strengthened as progressively more intervention data became available, reaching a Spearman correlation of 0.44 (p<0.001) using the first 50% of the intervention course (**Figure 2D**). We next evaluated whether these findings generalized to the independent PSOPT cohort. Using only the first 25% of intervention sessions, the locked facial signature achieved an external AUROC of 0.74 (95% CI: 0.52–0.92), with the corresponding FSS remaining positively associated with continuous EF improvement (Spearman ρ=0.35, p=0.031). Comparable performance was observed using the first 50% of intervention sessions (AUROC=0.79, 95% CI: 0.61–0.94; ρ=0.32, p=0.048) (**Figure 2E**; **Supplementary Table S5C**). Because interval estimates recompute the fixed feature definitions from window-level data, they derive from a different feature-construction pipeline than the primary analysis and are reported separately (**Supplementary Tables S5B** and **S5C**)

### Objective 3. Incremental value of the facial signature beyond baseline characteristics and conventional training measures

We next evaluated whether the facial signature provided information beyond participant characteristics available before treatment and conventional behavioral measures collected during intervention. In the BREATHE development cohort, conventional behavioral measures, including fatigue change, task accuracy, and IIV-RT, explained a moderate share of variance in subsequent EF improvement but discriminated responders near chance (base R²=0.23; AUROC=0.45; N=84). Adding FSS explained additional variance in EF improvement (ΔR²=0.193; p<0.001) and improved discrimination (ΔAUROC=0.33; 95% CI: 0.18–0.48; **Figure 3A**). These findings indicate that the facial signature captured information beyond conventional behavioral measures obtained during cognitive training.

**Figure 3.**
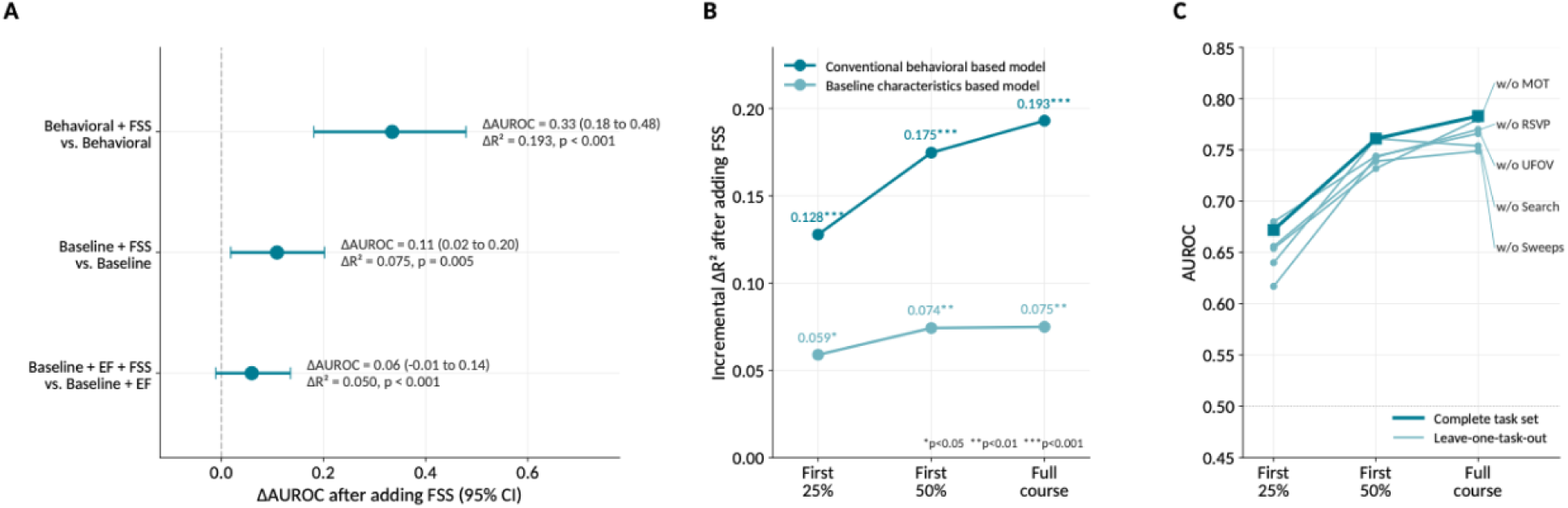
Incremental value and task robustness of the facial signature (BREATHE). (A) Change in AUROC (ΔAUROC) after adding the FSS to each base model, with 95% bootstrap confidence intervals from fixed-signature LOSO cross-validation. Annotations report the corresponding incremental explained variance (ΔR²) and one-degree-of-freedom partial-F-test (p) values. Because ΔAUROC was estimated from out-of-fold predictions whereas ΔR² was estimated in-sample, these metrics quantify different aspects of incremental value and may differ in statistical significance. (B) Incremental ΔR² after adding the FSS when the signature was reconstructed using the first 25%, first 50%, or complete intervention course (*p<0.05, **p<0.01, ***p<0.001). (C) Leave-one-task-out robustness across intervention-course intervals. Thin lines exclude each cognitive task in turn (labelled at right), the thick line uses the complete task set; dotted line indicates chance.

The FSS also provided incremental value beyond baseline participant characteristics. In models including age, sex, education, and ADSCT (N=85), adding the FSS increased explained variance (ΔR²=0.075; p=0.005) and improved discrimination (ΔAUROC=0.11; 95% CI: 0.02–0.20). After additional adjustment for baseline EF, the continuous contribution of the facial signature remained statistically significant (ΔR²=0.050; p<0.001), whereas the additional improvement in discrimination was modest and its confidence interval included zero (ΔAUROC=0.06, 95% CI: −0.01 to 0.14; **Figure 3A**).

We then evaluated whether the facial signature retained incremental value when computed from progressively shorter intervention-course intervals. Even when calculated from the first 25% of the intervention course, the FSS explained additional variance beyond both behavioral measures (ΔR²=0.128, p<0.001) and baseline characteristics (ΔR²=0.059, p=0.023). By the first 50% of the course, these increments increased to ΔR²=0.175 (p<0.001) and 0.074 (p=0.006), closely matching the full-course estimates of 0.193 and 0.075 (**Figure 3B**).

Finally, we examined whether the incremental association generalized to the independent PSOPT cohort. After adjustment for age, sex, education, and baseline EF (N=36), the incremental contribution of the FSS was directionally consistent with the development cohort but did not reach significance (ΔR²=0.027, partial-F p=0.31), reflecting the smaller external sample (**Supplementary Table S6**).

### Objective 4. Robustness of the facial signature

#### Across cognitive challenges

We first evaluated whether the facial signature depended on any individual therapeutic cognitive challenge included in the BREATHE intervention. In leave-one-task-out analyses, predictive performance remained similar after exclusion of each task, with AUROCs ranging from 0.75 to 0.78 compared with 0.78 using the complete task set. In contrast, prediction based on individual tasks alone showed greater variability ((AUROC=0.68–0.76; **Supplementary Figure S3A, Table S7A**), with the highest performance observed for MOT, followed by Search. A similar pattern was observed across intervention-course intervals. By the first 50% of the intervention course, leave-one-task-out AUROCs ranged from 0.73 to 0.76, compared with 0.76 for the complete task set (**Figure 3C; Supplementary Table S7A**), indicating that predictive performance remained stable even when individual tasks were excluded. Equivalent analyses could not be performed in PSOPT because task-level video recordings were unavailable.

#### Across intervention formats

We next evaluated whether the association between the facial signature and EF improvement differed across randomized intervention arms. In BREATHE, higher FSS was associated with greater baseline-adjusted EF improvement (β=0.30, p=0.002). However, there was no evidence that this association differed across intervention arms (FSS-by-arm interaction, p=0.38; **Supplementary Table S7B, Figure S3B**). Similar findings were observed in the MCI-only subgroup (interaction p=0.40) and within the mixed cognitive-status arm (FSS-by-cognitive-status interaction, ΔR²=0.003, p=0.62). In PSOPT, the arm-adjusted prognostic association was directionally positive but not statistically significant (β=0.33; p=0.083), with no significant FSS-by-arm interaction (p=0.46). Analyses using partial intervention-course data showed the same qualitative pattern but were underpowered to detect interaction effects.

## Discussion

In this study, we developed and externally validated a facial signature derived from passive facial dynamics recorded during therapeutic cognitive challenges to predict EF improvement following digital cognitive interventions in older adults. The facial signature generalized across two independent RCTs employing different cognitive intervention formats without feature reselection or outcome-informed model updating. Importantly, predictive performance was largely retained when the signature was reconstructed using only early intervention-course data, indicating that substantial predictive information is available before intervention completion. The facial signature also provided information beyond baseline participant characteristics and conventional behavioral measures collected during training, suggesting that it captures complementary intervention-process information not reflected by existing predictors. Finally, the signature remained robust across therapeutic cognitive challenges and randomized intervention arms, supporting its transportability across diverse digital cognitive intervention settings. Together, these findings identify passive facial dynamics during therapeutic cognitive challenge as a promising intervention-process marker of heterogeneity in EF improvement in older adults.

These findings suggest that response prediction for cognitive training should not be limited to static pretreatment variables. Baseline demographic, cognitive, clinical, and neuroimaging measures describe an individual’s predisposition before treatment begins, but they do not directly observe how that individual respond to repeated cognitive challenge. This intervention-process information may reflect individual differences in adaptation to repeated cognitive challenge, a behavioral dimension relevant to cognitive plasticity. The facial signature appeared to capture this intervention-process layer. It was not reducible to fatigue, task accuracy, or reaction-time variability, and it retained smaller but detectable information after adjustment for baseline EF. The early-session results show that a full-course-derived signature can be computed early and retain predictive information, but they do not establish that the signature could be discovered de novo using early-session data alone. Future prospective studies should therefore predefine the feature set, decision time point, and recalibration strategy before follow-up outcomes are available. Transparent reporting of development, validation, discrimination, and calibration is central for such prediction-model work.^26^

Mechanistically, the signature is most interpretable as a temporal facial-dynamics phenotype rather than as a set of isolated facial expressions. The AU10 and AU17 features reflected across-session instability in within-session lower-face dynamics, with greater instability associated with less EF improvement. The AU5 feature reflected typical upper-lid-raiser intensity across training sessions, with greater values associated with more EF improvement. This pattern should not be read as a simple emotion signal or as evidence that any single AU maps onto a single mental state. Instead, it suggests that the temporal organization and typical intensity of lower-face and eye-region behavior during repeated cognitive challenge may reflect individual differences in visual-attentional engagement, effort regulation, adaptation to task demand, or self-regulation. This interpretation is consistent with prior work showing that dynamic facial movements convey temporally structured, context-dependent information beyond static facial configurations and that facial cues can relate to attentional states during cognitively demanding tasks.^19,27,28^ However, these associations do not support a one-to-one mapping between any individual AU and a specific cognitive state.

The signature should be interpreted as a prognostic marker of likely improvement rather than as a treatment-selection marker. In the BEST biomarker framework, a predictive biomarker implies differential benefit from one treatment relative to another, whereas a prognostic biomarker indicates likely outcome independent of treatment assignment.^29^ Across randomized intervention arms, the FSS–EF association was positive, whereas FSS-by-arm interactions were not statistically supported. This pattern suggests that the signature may identify participants more likely to benefit across intervention conditions, but does not currently indicate which intervention a given participant should receive.

Clinically, the most plausible near-term use of the facial signature is low-burden monitoring and stratification rather than stand-alone decision-making. Because the marker is derived from ordinary training-session video, it could be embedded in remotely delivered cognitive interventions without additional sensors, clinic visits, or repeated self-report prompts. In future prospective studies, after a short early-training period, participants with lower predicted response could be evaluated for additional support, coaching, altered pacing, or extended training. In trials, the score could support enrichment or balancing of participants based on emerging response propensity. These applications differ from closed-loop intervention. The current model addresses “who may benefit?” but does not specify when to adapt, what to adapt, or how strongly to change the intervention. A closed-loop human–machine intervention would require a sensing layer, a control policy, and an actuation layer that changes task difficulty, pacing, novelty, content, or support. Facial dynamics could contribute to the sensing layer, but prospective adaptive trials are needed to show that acting on the signal improves outcomes.^4,30^

Several limitations should be considered. Both cohorts were modest in size, and the candidate feature space was large relative to the development sample. Although nested validation, permutation testing, and external validation reduced optimism, larger multi-site replications are needed. The responder definition captured any observed EF improvement rather than a psychometrically reliable or clinically meaningful threshold, and future studies should test reliable-change criteria, functional outcomes, and longer follow-up. The early-session analyses tested early deployment of a locked full-course-derived signature, not de novo early feature discovery. Automated AU extraction may be affected by lighting, camera angle, occlusion, eyewear, facial hair, skin tone, head motion, facial expressivity, and motor symptoms; broader verification and analytical validation under real-world home-use conditions are needed before deployment.^31,32^ Future work should combine facial AUs with ECG-derived heart-rate variability, pupillometry, eye tracking, task-level difficulty, subjective effort, fatigue, and neuroimaging to disentangle behavioral regulation, autonomic flexibility, and neural engagement. Prior work linking autonomic flexibility to learning and training-associated neuroplasticity in older adults suggests that ECG/HRV may be a particularly informative complementary modality.^33^

## Author contributions

Y.M.L. developed the analytical approach, analyzed the data, implemented the method, conducted the analyses, and drafted the manuscript. A.T. contributed to study design, and interpretation of the findings. M.A. contributed to data curation and preprocessing and validation of the analyses. K.L.H. contributed to the design and implementation of the BREATHE trial and data acquisition. C.T. contributed to the design and technical implementation of the PSOPT intervention and data acquisition. Y.Z. and E.A. contributed to computational methodology, computer-vision methodology, and analytical supervision. F.V.L. conceived and supervised the study, acquired funding, led the design and implementation of the clinical studies, interpreted the findings, and critically revised the manuscript. All authors reviewed and approved the final version of the manuscript.

## Acknowledgements

This study was funded by NIH AG081723, NR015452, and AG084471 and Stanford HAI seed funding. The funder played no role in study design, data collection, analysis and interpretation of data, or the writing of this manuscript.

## Data availability

The raw facial videos are not publicly available because they contain identifiable biometric information and are subject to participant-consent and institutional-review-board restrictions. Deidentified derived facial features, outcomes, and model predictions necessary to reproduce the reported analyses are available from the corresponding author upon reasonable request and subject to applicable institutional approvals and data-use agreements.

## Code availability

Analysis code used for facial-feature construction, FSS development, validation, and figure generation is available at FSS.

