## Supplementary materials for "Early-session facial dynamics predict executive-function response to digital cognitive interventions in older adults: development and external validation"

### Contents

Supplementary Methods 1. Cohort-specific cognitive challenges and measures

Supplementary Methods 2. Facial-data processing and multiscale feature construction

Supplementary Methods 3. Facial-signature development and model validation

Supplementary Methods 4. Secondary analyses, reproducibility, and data availability

Table S1. Cognitive challenges (A) and measurement definitions (B)

Table S2. Participant flow and analysis-specific sample sizes

Table S3. Complete multiscale facial-feature dictionary (2,208 features)

Table S4. Feature-screening cascade, AU-level ranking, and final signature

Table S5. Model specification, primary validation (A-B) and early-course interval analysis (C)

Table S6. Incremental-value analyses

Table S7. Task robustness and intervention heterogeneity

Figure S1. Participant flow and recording quality

Figure S2. Validation diagnostics

Figure S3. Exploratory robustness

### Supplementary Methods 1. Cohort-specific cognitive challenges and measures

#### Cognitive challenges

BREATHE delivered five processing-speed/attention (PS/A) cognitive challenges: multiple object tracking (MOT), rapid serial visual presentation (RSVP), visual search, visual sweep, and Useful Field of View (UFOV). PSOPT delivered four PS/A challenges: Motion of Target, Sound Sweeps, Mixed signal, and Delayed task switching. Task objectives, stimulus modality, participant responses and primary cognitive demands are given in Table S1A. The two programmes overlap in construct coverage (both include a multiple-object-tracking exercise targeting visuospatial working memory) but differ in composition: BREATHE was exclusively visual, whereas PSOPT additionally included auditory (Sound Sweeps) and multisensory (Mixed signal) exercises, and a task-switching exercise with a working-memory component (Delayed task switching). Task start and end times were coded for BREATHE recordings, enabling task-resolved facial analyses; PSOPT recordings were not task-labelled, so task-level analyses were possible only in BREATHE.

#### Outcome and covariate definitions

The executive-function (EF) composite was computed at pre- and post-intervention in each cohort as the mean of z-standardised scores from four executive tasks. In BREATHE (NIH EXAMINER) the components were dot counting, flanker, working memory and set shifting. In PSOPT (TabCAT) the components were DotCounting, Flanker, RunningDots and SetShifting, each standardised over all available participant-visits before averaging within participant-visit. The batteries share three components (dot counting, flanker, set shifting) and differ in the fourth, so composites are cohort-specific and are not on a common absolute scale; this motivated the cohort-independent responder definition and the use of rank-based statistics throughout.

EF change was defined as post- minus pre-intervention composite, and binary EF response as any measurable improvement (EF change > 0), applied identically in both cohorts. Conventional behavioral measures (BREATHE only) were mean within-session fatigue change, mean task accuracy, and intra-individual reaction-time variability (IIV-RT, the within-person coefficient of variation of reaction time). Mean reaction time was computed but deliberately excluded as a predictor because it was strongly correlated with baseline EF (Spearman rho = -0.64) and therefore behaves as a baseline-ability proxy rather than an intervention-process measure. Baseline characteristics were age, sex, years of education and Alzheimer's disease-signature cortical thickness (ADSCT), with baseline EF added in a sensitivity model. Full definitions and formulas are in Table S1B.

### Supplementary Methods 2. Facial-data processing and multiscale feature construction

Sessions were recorded with a workstation-mounted webcam, one video file per session. Native frame rate varied by recording: BREATHE (736 recordings) had a median of 29.8 frames per second with a mixed distribution (528 recordings at 28-32 fps, 61 at 25-28, 91 at 20-25 and 56 below 20 fps), whereas PSOPT (775 recordings) was near-uniform (773 at 28-32 fps; median 30.0 fps). Session duration was longer in BREATHE (median 55.2 minutes, IQR 44.5-58.9) than in PSOPT (median 41.0 minutes, IQR 40.1-47.4).

Videos were subsampled to 1 frame per second using a frame step of round(native fps / 1), so the step adapts to each file's native rate; each retained frame was stored with its exact timestamp. Faces were detected per frame with OpenCV YuNet (320x320 detection input, non-maximum-suppression threshold 0.3) with a Haar-cascade fallback; when multiple faces were present the largest was retained and cropped with padding. Frames without a detected face yielded no action-unit (AU) values and were excluded from all summaries, reducing the effective denominator rather than being imputed.

Face crops were encoded with py-feat 2.0.2 using the Detectorv1 (XGBoost AU) model; the v2 ConvNeXt AU model was evaluated as a sensitivity analysis and performed less well. Encoding produced 20 AUs (AU01, AU02, AU04, AU05, AU06, AU07, AU09, AU10, AU11, AU12, AU14, AU15, AU17, AU20, AU23, AU24, AU25, AU26, AU28, AU43) on a continuous 0-1 intensity scale, plus three head-pose variables (Pitch, Roll, Yaw).

Frames were aggregated into consecutive, non-overlapping 120-second analysis windows (stride equal to window length; median 119 frames per window, IQR 107-119, consistent with 1 fps sampling). A window was valid if at least one frame yielded AU output. Features were then constructed hierarchically. At the window level, each of the 23 channels was summarised by 2 statistics (mean and standard deviation across frames). At the session level, each window-level quantity was summarised by 12 statistics across the windows of that session: mean, standard deviation, median, interquartile range, minimum, maximum, range, within-session ordinary-least-squares slope across window order, first-tertile mean, last-tertile mean, last-minus-first-tertile change, and count of valid windows. Order-dependent statistics require at least three valid windows in a session. At the participant level, each session-level quantity was summarised by 4 aggregations across sessions: mean, standard deviation, median, and slope across session number.

This yields exactly 4 x 12 x 23 x 2 = 2,208 participant-level candidate features. Features are named training_<across-session aggregation>_<session-level statistic>_<channel>_<window-level statistic>; for example training_sd_early_late_change_AU10_mean is the across-session standard deviation of the within-session early-to-late change in the window-mean AU10 intensity. Quality control then removed all head-pose channels a priori and any AU channel whose window-level output was near-constant (robust 1st-99th-percentile range below 0.10), which flagged AU11 as saturated. Removing these four channels excluded 4 x 96 = 384 features, leaving 1,824 eligible AU features. Overall 80.7% of BREATHE windows were valid, giving a median of 155 valid windows per participant (IQR 118-175) across a median of 8 sessions; PSOPT gave a median of 359 valid windows (IQR 326-404) across a median of 18 sessions. The complete feature dictionary is provided as Table S3.

For the intervention-course interval analyses, the three selected feature definitions and their direction alignment were held fixed and recomputed from a participant-specific subset of sessions (the first 25%, first 50%, or the complete course, using a per-participant cutoff of ceil(fraction x that participant's maximum session number)). No feature discovery was repeated at any interval. Because order-dependent features require sufficient within- and between-session variation, the analysable sample declines at shorter intervals in BREATHE (Table S2).

### Supplementary Methods 3. Facial-signature development and model validation

Feature selection used BREATHE only, with continuous EF change as the development outcome, and the entire procedure was repeated within each cross-validation training fold. Candidate features were screened by their Spearman rank correlation with EF change at p < 0.05. A rank statistic was used because the EF-change outcome was strongly right-skewed (Shapiro-Wilk p < 10^-6) and most candidate features were non-normal, making a parametric screen unduly sensitive to outliers and to departures from linearity; a rank gate is also consistent with the robust statistic used for ranking. Pearson correlations were computed for the same features and are reported only as a sensitivity analysis. Development-stage p values were treated as feature filters, not as confirmatory inference.

Surviving features were deduplicated within AU, retaining per AU the feature with the largest absolute biweight midcorrelation with EF change; the biweight midcorrelation is a robust correlation that down-weights outlying observations via a median/median-absolute-deviation weighting. The resulting AU-level features were ranked by the same robust statistic and the top three were taken, subject to a prespecified directional-coverage rule requiring that both positive and negative associations be represented (if the top three were single-signed, the weakest was replaced by the highest-ranked feature of the missing direction). Signature size was fixed a priori at K = 3 by an events-per-variable criterion of at least 10 outcome events per predictor (54 responders / 3 = 18). The full cascade is given in Table S4A: 2208 candidates, 1824 after quality control, 187 after the Spearman screen, 17 after within-AU deduplication, and 3 final features.

The final signature comprises training_sd_early_late_change_AU10_mean (across-session standard deviation of the within-session early-to-late change in window-mean AU10; negative direction), training_sd_session_sd_AU17_mean (across-session standard deviation of the within-session standard deviation of window-mean AU17; negative direction), and training_median_session_median_AU05_mean (across-session median of the within-session median of window-mean AU05; positive direction). The equal-weight Facial Signature Score was defined as FSS = [-z(AU10 feature) - z(AU17 feature) + z(AU05 feature)] / 3, and was evaluated separately from the fitted model as an interpretable summary.

Binary EF response was predicted directly by the equal-weight FSS. Because the FSS applies fixed unit weights to the three sign-aligned standardised features, no feature weights were fitted, and discrimination was therefore the AUROC of the FSS itself. To place predictions on a probability scale for calibration, a single univariable logistic link of EF response on the FSS was fitted in BREATHE and then locked: logit(p) = 0.61 + 1.60 x FSS. Because this link is strictly monotone, AUROC is identical on the FSS and probability scales. The nominal operating threshold was 0.50 predicted probability, set in BREATHE before external validation. No hyperparameter search was performed. Standardisation parameters were estimated on training data only. Continuous EF change was additionally modelled by ridge regression (alpha = 1.0) as a secondary analysis, not reported in the main text. The locked FSS formula, probability link and development scaling parameters are given in Table S5A.

Internal performance was estimated by fully nested leave-one-subject-out (LOSO) cross-validation, in which screening, deduplication, ranking, standardisation and model fitting were all repeated using only the training participants of each fold, so that the held-out participant influenced no step. A fixed-signature LOSO estimate, in which the three features were fixed on the full cohort and only standardisation and fitting were repeated per fold, is reported as a deliberately more optimistic reference. Feature-selection stability was quantified as the proportion of outer folds in which each AU was reselected. To quantify optimism arising from feature search, the complete development pipeline including feature selection was repeated across 1,000 participant-level outcome permutations. Confidence intervals were obtained by participant-level bootstrap resampling with 2,000 replicates; for cross-validated metrics participants and their out-of-fold predictions were resampled together, and for paired model comparisons both models' predictions were resampled together.

External validation applied the locked BREATHE model to PSOPT with no feature reselection, no refitting of coefficients and no threshold modification. In the primary frozen analysis the development standardisation parameters were also applied unchanged. Because the cohorts differed in participant characteristics, intervention delivery and recording conditions, a prespecified outcome-independent sensitivity analysis re-estimated the feature-standardisation parameters from the PSOPT predictor distribution alone; feature definitions, model coefficients and the classification threshold remained locked, and PSOPT outcomes were never used. Discrimination was summarised by AUROC, calibration by calibration slope, calibration intercept and Brier score, and external discrimination was additionally tested by a conditional permutation test (1,000 permutations of PSOPT outcome labels with all model parameters fixed). Results appear in Table S5B and Figure S2.

### Supplementary Methods 4. Secondary analyses, reproducibility, and data availability

Intervention-course intervals. IMPORTANT: the interval analysis is a distinct pipeline from the primary analysis. Primary estimates (Table S5B) use the precomputed participant-level feature table (BREATHE n = 87), whereas interval estimates (Table S5C) recompute the three fixed feature definitions from window-level data at each interval (BREATHE n = 88 at full course). Full-course interval estimates are therefore close to, but not identical with, the primary estimates and must not be reported as the same quantity.

Performance was evaluated at the first 25%, first 50% and complete course. In BREATHE, interval-specific feature-standardisation parameters were estimated from the corresponding BREATHE data and the same equal-weight scoring rule was applied, with nothing refitted; in PSOPT the matched interval was evaluated under the same outcome-independent recalibration strategy used for the primary external analysis. The selected features and their direction alignment were fixed across all intervals.

Incremental value. Three interpretable base models were specified in BREATHE: a behavioral model (mean within-session fatigue change, mean task accuracy, IIV-RT), a baseline-characteristics model (age, sex, education, ADSCT), and a sensitivity model additionally including baseline EF. Incremental continuous value was quantified as the change in R-squared with a one-degree-of-freedom partial F test, computed in-sample on continuous EF change. Incremental classification value was quantified as the change in fixed-signature LOSO AUROC with a paired bootstrap 95% confidence interval. These two quantities are distinct estimands: the former is an in-sample fit statistic for a continuous outcome, the latter a cross-validated discrimination statistic for a binary outcome, and they are therefore not expected to agree in statistical significance. In PSOPT the incremental analysis was adjusted for age, sex, education and baseline EF only, because ADSCT and the behavioral measures were unavailable externally. Results appear in Table S6.

Task robustness. In BREATHE the fixed three-feature signature was recomputed (i) excluding one cognitive challenge at a time and (ii) from each challenge alone, in each case rebuilding the participant-level features from only the retained windows, and re-evaluated by fixed-signature LOSO AUROC and continuous LOSO regression. Equivalent analyses were not possible in PSOPT because recordings were not task-labelled. Results appear in Table S7A and Figure S3A.

Intervention heterogeneity. In each randomised cohort, post-intervention EF was modelled as a function of baseline EF, the FSS, intervention arm and the FSS-by-arm interaction; the omnibus interaction was the treatment-heterogeneity test and the FSS main effect the prognostic estimate, reported as a fully standardised coefficient. The interaction was repeated in the MCI-only subgroup, and an FSS-by-cognitive-status interaction was tested within the CCT+RFB arm, the only arm enrolling both cognitively unimpaired and MCI participants. All arm models were fitted by ordinary least squares with heteroskedasticity-consistent (HC3) robust standard errors, which remain valid when residual variance is not constant across participants. These analyses are exploratory and underpowered for interaction effects. Results appear in Table S7B and Figure S3B.

Statistical reporting. Discrimination was summarised by AUROC and continuous associations by Spearman rank correlations, reflecting use of the FSS as a response-ranking marker. Empirical one-sided permutation p values were computed as p = (b + 1) / (B + 1), where b is the number of permuted statistics at least as large as the observed statistic and B the number of permutations; these tests are one-sided because the alternative hypothesis is directional (discrimination better than chance), so only upper-tail exceedances constitute evidence. Other tests were two-sided at a nominal level of 0.05.

Software and reproducibility. Analyses used Python 3.11.15 with scikit-learn 1.7.2, statsmodels 0.14.6, SciPy 1.11.4, NumPy 1.26.4, pandas 2.3.3 and py-feat 2.0.2, with random seed 42. The execution order was: video metadata and frame sampling; face detection and cropping; py-feat encoding; 120-second window alignment; session-level and participant-level feature construction; feature screening and signature development with nested cross-validation; locked external validation and recalibration; then the secondary interval, incremental, task-robustness and heterogeneity analyses.

Data availability. Derived participant-level facial features, the fitted signature specification and the analysis code required to reproduce every table and figure in this Supplement can be made available to qualified investigators under a data-use agreement. Raw facial video cannot be shared: it is inherently identifiable biometric data, and consent was not obtained for redistribution. Requests for derived data should be directed to the corresponding author and will be considered subject to institutional review board approval and a data-use agreement.

### Table S1A. Cohort-specific cognitive challenges

Table S1A. Cognitive challenges delivered in each cohort. Descriptions are taken from the study task documentation. Task-specific video labels indicate whether facial recordings could be attributed to individual tasks, which determined whether task-resolved analyses were possible. Abbreviations: MOT, multiple object tracking; RSVP, rapid serial visual presentation; UFOV, Useful Field of View.

| **cohort** | **task** | **procedure** | **modality** | **demands** | **response** | **outputs** | **video labels** |
| --- | --- | --- | --- | --- | --- | --- | --- |
| BREATHE | Multiple object tracking (MOT) | Participants track a varying number of target objects, defined by their spatiotemporal locations, among visually similar distractors. | Visual | Visuospatial working memory capacity | Identify the tracked target objects | Accuracy; reaction time; intra-individual RT variability | Yes |
| BREATHE | Rapid serial visual presentation (RSVP) | Participants view a stream of images in rapid succession and determine whether a target image appears, as quickly as possible. | Visual | Visual processing speed | Speeded target-present detection | Accuracy; reaction time; intra-individual RT variability | Yes |
| BREATHE | Visual search | Participants locate a target stimulus among a field of distractor stimuli as quickly as possible. | Visual | Visual attentional processing | Speeded target localisation | Accuracy; reaction time; intra-individual RT variability | Yes |
| BREATHE | Visual sweep | Participants observe a spatial-frequency sweep (movement of bars) and determine whether the sweep moves inward or outward. | Visual | Visual processing speed | Two-alternative forced choice (inward vs outward) | Accuracy; reaction time; intra-individual RT variability | Yes |
| BREATHE | Useful Field of View (UFOV) | Measures the ability to extract information in a single glance without moving the head or eyes. Participants are tested on central visual processing speed, divided attention between central and peripheral stimuli, and selective attention to peripheral stimuli among distractors. | Visual | Attentional processing (central, divided, selective) | Central and peripheral stimulus identification/localisation | Accuracy; reaction time; intra-individual RT variability, recorded separately for central and peripheral conditions | Yes |
| PSOPT | Motion of Target | Multiple-object-tracking exercise in which participants track a varying number of target objects, defined by their spatio-temporal location, among visually identical distractors. | Visual | Visuo-spatial working memory capacity | Identify the tracked target objects | - | No - PSOPT recordings were not task-labelled |
| PSOPT | Sound Sweeps | Time-order-judgement exercise in which two successive frequency-modulated tone sweeps are presented and participants indicate whether the pitch (frequency) increased or decreased within each tone. | Auditory | Auditory processing speed | Two-alternative forced choice (rising vs falling pitch) for each sweep | - | No - PSOPT recordings were not task-labelled |
| PSOPT | Mixed signal | Stroop-based exercise requiring participants to attend to target information while ignoring competing information; participants listen to a letter, colour, number or other item while simultaneously viewing a set of letters, symbols, numbers or words. | Audio-visual (multisensory) | Multisensory attention and cognitive control (interference resolution) | Respond to the attended modality while ignoring the competing stream | - | No - PSOPT recordings were not task-labelled |
| PSOPT | Delayed task switching | Basic task-switching design in which participants view a target stimulus (indicated by an arrow) and two options, and select the better match to the target according to a specified criterion (e.g. colour or shape). The criterion varies from trial to trial and is presented a few seconds before the trial but is not shown on screen during the trial, so it must be held in working memory. | Visual | Task switching / cognitive flexibility and working memory | Two-alternative forced choice (select the matching option) | - | No - PSOPT recordings were not task-labelled |

### Table S1B. Measurement definitions

Table S1B. Operational definitions of outcomes and covariates. EF, executive function; IIV-RT, intra-individual reaction-time variability; ADSCT, Alzheimer's disease-signature cortical thickness; FSS, Facial Signature Score.

| **measure** | **definition** | **formula** | **aggregation** | **cohorts** | **use** |
| --- | --- | --- | --- | --- | --- |
| EF composite | Executive-function composite computed at pre- and post-intervention as the mean of z-standardised scores from four executive tasks | BREATHE (NIH EXAMINER): mean z of dot counting (dc), flanker (fl), working memory (wm) and set shifting (ss) [label/breathe_examiner_both_sites.csv: dc_score, fl_score, wm_score, ss_score -> dc_z..ss_z -> cog_composite_4task]. PSOPT (TabCAT): mean z of DotCounting_TotalScore, Flanker_TotalScore, RunningDots_TrialScore and SetShifting_TotalScore, each standardised over all available participant-visits before averaging within participant-visit [label/calculate_ef_composite_4task.R] | Task -> participant x timepoint | Both (cohort-specific batteries) | Primary outcome source; cohort-specific and therefore not on a common absolute scale |
| EF change | Post-intervention minus pre-intervention EF composite | ef_change = ef_post - ef_pre | Participant | Both | Development outcome for feature screening; continuous secondary outcome |
| Binary EF response | Any measurable improvement in EF | ef_responder = 1 if ef_change > 0, else 0 | Participant | Both | Primary outcome for classification (AUROC); cohort-independent definition |
| Within-session fatigue change | Change in self-reported fatigue from before to after a training session (visual analogue scale) | fatigue_change = fatigue_mean_post - fatigue_mean_pre per session, averaged across sessions (fat_mean) | Session -> participant | BREATHE only | Behavioral incremental model |
| Mean task accuracy | Mean accuracy across the five BREATHE training tasks | {task}_accuracy z-scored across participants, averaged across tasks within session, then averaged across sessions (acc_mean) | Task -> session -> participant | BREATHE only | Behavioral incremental model |
| IIV-RT (intra-individual reaction-time variability) | Within-person variability of reaction time, expressed as the coefficient of variation | {task}_iivrt_cv = SD(RT)/mean(RT) within task and session; z-scored across participants, averaged across tasks, then across sessions (iiv_mean) | Task -> session -> participant | BREATHE only | Behavioral incremental model |
| Mean reaction time (excluded from models) | Mean reaction time across training tasks | {task}_mean_rt aggregated as above (rt_mean) | Task -> session -> participant | BREATHE only | NOT used as a predictor: strongly correlated with baseline EF (Spearman rho = -0.64), so it behaves as a baseline-ability proxy rather than an intervention-process measure |
| ADSCT (Alzheimer's disease-signature cortical thickness) | Structural-MRI cortical-thickness index over an Alzheimer's-disease signature region set, with lower values indicating greater neurodegeneration | Project variable adsct_jack.1 (label/BREATHE_merged.csv) | Participant (baseline) | BREATHE only (unavailable in PSOPT) | Baseline-characteristics model |
| Baseline EF | Pre-intervention EF composite | ef_pre | Participant (baseline) | Both | Covariate in the baseline-EF sensitivity model and the baseline term in all ANCOVA arm models |

### Table S2. Participant flow and analysis-specific sample sizes

Table S2. Participant flow and the analysable sample for every reported analysis. Sample sizes differ across analyses because completeness requirements differ (for example, arm models require pre- and post-intervention EF rather than the facial signature, and interval analyses require that order-dependent features be computable from the restricted session set). Counts are participants.

| **cohort** | **analysis** | **interval or subgroup** | **initial eligible N** | **excluded N** | **final N** | **exclusion reason** | **required variables** |
| --- | --- | --- | --- | --- | --- | --- | --- |
| BREATHE | Source records | - | 123 | 20 | 103 | No usable session video | facial signature |
| BREATHE | Outcome linkage | - | 103 | 13 | 90 | No linked pre/post EF assessment | EF pre/post |
| BREATHE | EF change computable | - | 90 | 2 | 88 | Responder label present but EF change not computable | ef_change |
| BREATHE | Primary nested LOSO model | full course | 88 | 1 | 87 | Incomplete AU10 feature | 3 features + ef_change/responder |
| BREATHE | Fixed-signature LOSO | full course | 88 | 1 | 87 | Incomplete AU10 feature | 3 features + ef_responder |
| BREATHE | FSS-EF association | full course | 88 | 1 | 87 | Incomplete AU10 feature | 3 features + ef_change |
| BREATHE | Behavioral incremental model | full course | 87 | 3 | 84 | Missing fatigue/accuracy/IIV-RT | 3 features + ef_change + fatigue,accuracy,IIV-RT |
| BREATHE | Behavioral incremental model | first 50% | 82 | 4 | 78 | Missing behavioral covariates or non-computable interval features | as above (interval-rebuilt) |
| BREATHE | Behavioral incremental model | first 25% | 71 | 3 | 68 | Missing behavioral covariates or non-computable interval features | as above (interval-rebuilt) |
| BREATHE | Baseline-characteristics model | full course | 87 | 2 | 85 | Missing age/sex/education/ADSCT | 3 features + ef_change + age,sex,education,ADSCT |
| BREATHE | Baseline-characteristics model | first 50% | 82 | 1 | 81 | Missing covariates or non-computable interval features | as above (interval-rebuilt) |
| BREATHE | Baseline-characteristics model | first 25% | 71 | 1 | 70 | Missing covariates or non-computable interval features | as above (interval-rebuilt) |
| BREATHE | Baseline-EF sensitivity model | full course | 87 | 2 | 85 | Missing covariates or baseline EF | baseline covariates + ef_pre |
| BREATHE | Interval signature analysis | first 25% | 88 | 17 | 71 | Interval features not computable (too few sessions/windows) | 3 interval-rebuilt features + EF |
| BREATHE | Interval signature analysis | first 50% | 88 | 6 | 82 | Interval features not computable | 3 interval-rebuilt features + EF |
| BREATHE | Interval signature analysis | full course | 88 | 0 | 88 | - | 3 interval-rebuilt features + EF |
| BREATHE | Leave-one-task-out | full course | 88 | 0 | 88 | - | task-restricted rebuilt features + EF |
| BREATHE | Single-task only: MOT | full course | 88 | 0 | 88 | - | task-restricted rebuilt features + EF |
| BREATHE | Single-task only: RSVP | full course | 88 | 0 | 88 | - | task-restricted rebuilt features + EF |
| BREATHE | Single-task only: Search | full course | 88 | 1 | 87 | Features not computable from this task alone | task-restricted rebuilt features + EF |
| BREATHE | Single-task only: Sweeps | full course | 88 | 9 | 79 | Features not computable from this task alone | task-restricted rebuilt features + EF |
| BREATHE | Single-task only: UFOV | full course | 88 | 14 | 74 | Features not computable from this task alone | task-restricted rebuilt features + EF |
| BREATHE | Arm model, all participants | full course | 88 | 0 | 88 | - | ef_pre, ef_post, FSS, randomisation |
| BREATHE | Arm model, MCI only | full course | 88 | 28 | 60 | Cognitively unimpaired participants excluded | as above, MCI subset |
| BREATHE | CCT+RFB cognitive-status model | full course | 51 | 0 | 51 | - | as above, arm 1 only (28 CU, 23 MCI) |
| PSOPT | Source records | - | 54 | 10 | 44 | No session video available | facial signature |
| PSOPT | Usable recording + EF change | - | 44 | 4 | 40 | Recording quality criteria unmet or missing post EF | usable_strict + ef_change |
| PSOPT | Frozen external validation | full course | 40 | 1 | 39 | Incomplete AU10/AU17 feature | 3 features + ef_responder/ef_change |
| PSOPT | Recalibrated external validation | full course | 40 | 1 | 39 | Incomplete AU10/AU17 feature | 3 features + ef_responder/ef_change |
| PSOPT | Interval validation | first 25% | 40 | 1 | 39 | Incomplete features | 3 interval-rebuilt features + EF |
| PSOPT | Interval validation | first 50% | 40 | 1 | 39 | Incomplete features | 3 interval-rebuilt features + EF |
| PSOPT | Adjusted incremental model | full course | 39 | 3 | 36 | Missing age/sex/education/baseline EF | 3 features + ef_change + age,sex,education,ef_pre |
| PSOPT | Arm model | full course | 40 | 0 | 40 | - | ef_change, FSS, randomisation group (21 vs 19) |

### Table S3. Complete multiscale facial-feature dictionary

Table S3. All 2,208 participant-level candidate features, formed as 4 across-session aggregations x 12 session-level statistics x 23 channels (20 action units and 3 head-pose variables) x 2 window-level statistics. QC status indicates eligibility for feature screening: head-pose channels were excluded a priori and AU11 was excluded as near-constant (detector saturation), removing 384 features and leaving 1,824 eligible. Features are named training_<across-session aggregation>_<session-level statistic>_<channel>_<window-level statistic>.

**See “supplementary_table_S3_npj-aging.xlsx” for the full Table S3.**

### Table S4A. Feature-screening cascade

Table S4A. Sequential reduction from candidate features to the final three-feature signature, performed in BREATHE only. Screening p values are development-stage filters, not confirmatory tests.

| **stage** | **n entering** | **n retained** | **criterion** |
| --- | --- | --- | --- |
| 1. Candidate multiscale features | - | 2208 | 4 across-session aggregations x 12 session-level statistics x 23 channels x 2 window-level statistics |
| 2. Channel-level quality control | 2208 | 1824 | remove head-pose channels (Pitch/Roll/Yaw) and near-constant AU channels (AU11) |
| 3. Spearman screen vs continuous EF change | 1824 | 187 | Spearman rank correlation p < 0.05 (development-stage filter, not confirmatory) |
| 4. Within-AU deduplication | 187 | 17 | retain, per AU, the feature with the largest \|biweight midcorrelation\| |
| 5. Final signature | 17 | 3 | top-K by \|biweight midcorrelation\| with K=3 fixed a priori (EPV>=10) and bidirectional-coverage constraint |

### Table S4B. AU-level screening and ranking

Table S4B. All AU-level candidates remaining after within-AU deduplication, ranked by absolute biweight midcorrelation with continuous EF change in BREATHE. Spearman rho and the screening p value are development-stage statistics. Direction refers to the sign of the association with EF change.

| **rank** | **AU** | **retained feature name** | **spearman rho** | **screening p** | **abs biweight midcorrelation** | **association direction** | **final FSS inclusion** |
| --- | --- | --- | --- | --- | --- | --- | --- |
| 1 | AU10 | training_sd_early_late_change_AU10_mean | -0.3572938689217759 | 0.0006805797827668 | 0.3752644405087917 | negative | yes |
| 2 | AU17 | training_sd_session_sd_AU17_mean | -0.3985770138424149 | 0.0001199719873841 | 0.3546900780683745 | negative | yes |
| 3 | AU20 | training_median_session_sd_AU20_mean | -0.3593216160050721 | 0.000585763765098 | 0.3253787570534506 | negative | no |
| 4 | AU05 | training_median_session_median_AU05_mean | 0.3082138705927935 | 0.0034841925107789 | 0.3236490434651594 | positive | yes |
| 5 | AU06 | training_median_session_iqr_AU06_mean | -0.3302983339790779 | 0.0016729754970587 | 0.3197328537208143 | negative | no |
| 6 | AU12 | training_median_session_median_AU12_mean | -0.3144482406396393 | 0.002847999295566 | 0.3193751096365367 | negative | no |
| 7 | AU14 | training_sd_session_min_AU14_mean | -0.3339086330175055 | 0.0014761623772922 | 0.2801243074099845 | negative | no |
| 8 | AU04 | training_slope_session_min_AU04_std | 0.2295974076291783 | 0.0314088648897208 | 0.2770007323943694 | positive | no |
| 9 | AU15 | training_sd_late_mean_AU15_std | -0.298281145433412 | 0.0047624797563157 | 0.2556638602862736 | negative | no |
| 10 | AU23 | training_sd_early_late_change_AU23_mean | -0.2379529051541881 | 0.0264624405756797 | 0.2547075699361203 | negative | no |
| 11 | AU01 | training_sd_within_session_slope_AU01_std | -0.2811982670564616 | 0.0079559022940817 | 0.2533872088731703 | negative | no |
| 12 | AU07 | training_sd_within_session_slope_AU07_mean | -0.2155964918460076 | 0.0436592409763348 | 0.2464972697717517 | negative | no |
| 13 | AU25 | training_sd_early_late_change_AU25_mean | -0.2240832543559087 | 0.0369341587934223 | 0.2442984008819146 | negative | no |
| 14 | AU28 | training_slope_early_late_change_AU28_std | 0.3034737916454035 | 0.0042714265866754 | 0.2370461978798479 | positive | no |
| 15 | AU43 | training_sd_early_late_change_AU43_mean | -0.2152074068673908 | 0.0453052093234883 | 0.2263004277127789 | negative | no |
| 16 | AU24 | training_slope_session_iqr_AU24_std | 0.2452713888204008 | 0.0212645987238512 | 0.2218503059126503 | positive | no |
| 17 | AU02 | training_slope_session_min_AU02_mean | -0.2175337254763834 | 0.0417584508107216 | 0.1776359650230191 | negative | no |

### Table S4C. Final three-feature signature

Table S4C. The locked signature. FSS sign is the direction alignment applied when constructing the equal-weight Facial Signature Score. Selection frequency is the proportion of nested leave-one-subject-out folds (87 folds) in which the corresponding action unit was reselected by the full development pipeline. Distribution summary is the median (interquartile range) in the BREATHE development cohort.

| **feature name** | **AU** | **plain language definition** | **FSS sign** | **spearman rho** | **abs biweight midcorrelation** | **nested LOSO AU selection frequency** | **distribution median IQR** | **n nonmissing** |
| --- | --- | --- | --- | --- | --- | --- | --- | --- |
| training_sd_early_late_change_AU10_mean | AU10 | Across-session SD of the within-session early-to-late change in window-mean AU10 intensity | -1 | -0.357 | 0.375 | 99% | 0.077 (0.050-0.100) | 89 |
| training_sd_session_sd_AU17_mean | AU17 | Across-session SD of the within-session SD of window-mean AU17 intensity | -1 | -0.399 | 0.355 | 100% | 0.010 (0.007-0.014) | 90 |
| training_median_session_median_AU05_mean | AU05 | Across-session median of the within-session median of window-mean AU05 intensity | 1 | 0.308 | 0.324 | 97% | 0.388 (0.364-0.410) | 90 |

### Table S5A. Locked model specification

Table S5A. Specification of the locked marker of record. The FSS applies fixed unit weights to the three sign-aligned standardised features, so no feature weights were fitted; only the univariable probability link has estimated parameters. The FSS formula, the link, the threshold and the development scaling parameters were never re-estimated in PSOPT.

| **item** | **value** |
| --- | --- |
| Marker of record | Equal-weight Facial Signature Score (FSS); no fitted feature weights |
| FSS formula | FSS = [-z(training_sd_early_late_change_AU10_mean) - z(training_sd_session_sd_AU17_mean) + z(training_median_session_median_AU05_mean)] / 3 |
| Probability link (locked, development-fitted) | logit(p) = +0.6126 +1.6002 x FSS |
| Discrimination | AUROC of the FSS; the link is monotone so AUROC is identical on the probability scale |
| Standardisation | z-scoring of each feature; parameters estimated on development data only (frozen) or re-estimated from the external predictor distribution alone (recalibration sensitivity analysis) |
| Nominal operating threshold | 0.50 (predicted probability) |
| Secondary continuous model | Ridge regression (alpha = 1.0) on z-scored EF change, retained as a secondary predictive analysis |
| Development scaling (mean, SD): training_sd_early_late_change_AU10_mean | 0.0767, 0.0370 |
| Development scaling (mean, SD): training_sd_session_sd_AU17_mean | 0.0110, 0.0058 |
| Development scaling (mean, SD): training_median_session_median_AU05_mean | 0.3890, 0.0369 |

### Table S5B. Primary validation, calibration and permutation

Table S5B. Primary validation. Nested estimates re-run the entire development pipeline within each training fold; fixed-signature estimates hold the three features fixed and are therefore optimistic. Frozen applies the development standardisation unchanged; recalibrated re-estimates standardisation parameters from the PSOPT predictor distribution only, leaving features, coefficients and threshold locked. All estimates use the precomputed participant-level feature table (the primary pipeline); interval estimates from the window-rebuilt pipeline are reported separately in Table S5C and are not the same quantity. Confidence intervals are participant-level bootstrap percentile intervals (2,000 replicates). Spearman rho is the association between the FSS and continuous EF change. Calibration for the BREATHE fixed-signature row is in-sample by construction (slope 1.00, intercept 0.00).

| **cohort analysis** | **n** | **model status** | **auroc** | **ci95** | **spearman rho** | **spearman p** | **brier** | **calib slope** | **calib intercept** |
| --- | --- | --- | --- | --- | --- | --- | --- | --- | --- |
| BREATHE nested LOSO (FSS) | 87 | features, direction alignment and scaling re-derived within each fold | 0.768 | 0.66-0.86 | 0.419 | 5.34e-05 |  |  |  |
| BREATHE fixed-signature (FSS) | 87 | features fixed on the full development cohort; development scaling | 0.796 | 0.70-0.89 | 0.488 | 1.67e-06 | 0.182 | 1.00 (by construction) | 0.00 (by construction) |
| PSOPT frozen (FSS) | 39 | locked FSS formula + locked link; frozen standardisation | 0.781 | 0.60-0.93 | 0.303 | 0.06 | 0.248 | 0.71 | 1.33 |
| PSOPT recalibrated (FSS) | 39 | locked FSS formula + locked link; standardisation re-estimated (outcome-independent) | 0.79 | 0.60-0.94 | 0.323 | 0.045 | 0.167 | 1.19 | 0.27 |

### Table S5C. Early-course performance (interval analysis)

Table S5C. Performance across intervention-course intervals. This is the INTERVAL pipeline: the three fixed feature definitions are recomputed from window-level data for each interval, so full-course values here differ slightly from the primary estimates in Table S5B (different feature-construction pipeline and, in BREATHE, a different analysable sample: n = 88 versus n = 87). PSOPT intervals use outcome-independent recalibration throughout, per the prespecified interval strategy; they are therefore not the frozen external result. Confidence intervals are participant-level bootstrap percentile intervals (2,000 replicates).

| **cohort** | **interval** | **pipeline** | **scaling** | **auroc** | **ci95** | **spearman rho** | **p** |
| --- | --- | --- | --- | --- | --- | --- | --- |
| BREATHE | first 25% | features rebuilt from window-level data (interval analysis) | internal (development) | 0.672 | 0.54-0.79 | 0.307 | 0.009 |
| BREATHE | first 50% | features rebuilt from window-level data (interval analysis) | internal (development) | 0.761 | 0.65-0.86 | 0.439 | <0.001 |
| BREATHE | full course | features rebuilt from window-level data (interval analysis) | internal (development) | 0.783 | 0.69-0.88 | 0.47 | <0.001 |
| PSOPT | first 25% | features rebuilt from window-level data (interval analysis) | outcome-independent recalibration | 0.741 | 0.52-0.92 | 0.346 | 0.031 |
| PSOPT | first 50% | features rebuilt from window-level data (interval analysis) | outcome-independent recalibration | 0.793 | 0.61-0.94 | 0.319 | 0.048 |
| PSOPT | full course | features rebuilt from window-level data (interval analysis) | outcome-independent recalibration | 0.79 | 0.60-0.94 | 0.323 | 0.045 |

### Table S6. Incremental-value analyses

Table S6. Incremental value of the FSS over interpretable base models. Delta R-squared and the one-degree-of-freedom partial F test are in-sample estimates for continuous EF change; delta AUROC is cross-validated (fixed-signature leave-one-subject-out) for binary EF response with a paired bootstrap 95% confidence interval. These are different estimands and are not expected to agree in significance.

| **cohort** | **interval** | **base model** | **covariates** | **n** | **base R2** | **R2 with FSS** | **delta R2** | **partial F** | **partial F p** | **base auroc** | **auroc with FSS** | **delta auroc** | **delta auroc ci95** |
| --- | --- | --- | --- | --- | --- | --- | --- | --- | --- | --- | --- | --- | --- |
| BREATHE | first 25% | Behavioral | mean within-session fatigue change, mean task accuracy, IIV-RT | 68 | 0.295 | 0.423 | 0.128 | 13.96 | <0.001 | 0.456 | 0.61 | 0.154 | -0.00 to 0.31 |
| BREATHE | first 25% | Baseline characteristics | age, sex, education years, ADSCT | 70 | 0.247 | 0.306 | 0.059 | 5.43 | 0.023 | 0.744 | 0.797 | 0.053 | -0.01 to 0.12 |
| BREATHE | first 25% | Baseline characteristics + baseline EF | age, sex, education years, ADSCT, baseline EF | 70 | 0.66 | 0.728 | 0.068 | 15.71 | <0.001 | 0.809 | 0.848 | 0.04 | -0.03 to 0.11 |
| BREATHE | first 50% | Behavioral | mean within-session fatigue change, mean task accuracy, IIV-RT | 78 | 0.228 | 0.403 | 0.175 | 21.35 | <0.001 | 0.432 | 0.768 | 0.336 | 0.18 to 0.49 |
| BREATHE | first 50% | Baseline characteristics | age, sex, education years, ADSCT | 81 | 0.228 | 0.302 | 0.074 | 7.98 | 0.006 | 0.741 | 0.839 | 0.097 | 0.02 to 0.19 |
| BREATHE | first 50% | Baseline characteristics + baseline EF | age, sex, education years, ADSCT, baseline EF | 81 | 0.642 | 0.703 | 0.061 | 15.21 | <0.001 | 0.821 | 0.889 | 0.068 | 0.00 to 0.15 |
| BREATHE | full course | Behavioral | mean within-session fatigue change, mean task accuracy, IIV-RT | 84 | 0.228 | 0.421 | 0.193 | 26.32 | <0.001 | 0.447 | 0.782 | 0.335 | 0.18 to 0.48 |
| BREATHE | full course | Baseline characteristics | age, sex, education years, ADSCT | 85 | 0.202 | 0.277 | 0.075 | 8.19 | 0.005 | 0.699 | 0.808 | 0.109 | 0.02 to 0.20 |
| BREATHE | full course | Baseline characteristics + baseline EF | age, sex, education years, ADSCT, baseline EF | 85 | 0.639 | 0.689 | 0.05 | 12.57 | <0.001 | 0.81 | 0.869 | 0.06 | -0.01 to 0.14 |
| PSOPT | full course | Baseline characteristics + baseline EF (external) | age, sex, education, baseline EF (ADSCT and behavioral measures unavailable) | 36 | 0.226 | 0.253 | 0.027 | [not stored] | 0.305 | 0.696 | 0.742 | 0.046 | [not stored; exploratory] |

### Table S7A. Task robustness

Table S7A. Robustness of the fixed three-feature signature to the composition of the training programme in BREATHE, evaluated by fixed-signature leave-one-subject-out AUROC. Continuous-association estimates are Spearman correlations between the leave-one-subject-out continuous prediction and EF change. Confidence intervals are participant-level bootstrap percentile intervals (2,000 replicates, seed 42). These analyses are exploratory sensitivity analyses and the intervals are wide, particularly for single-task reconstructions where fewer participants have computable features. Equivalent analyses were not possible in PSOPT because recordings were not task-labelled.

| **analysis type** | **task included or excluded** | **interval** | **n** | **auroc** | **auroc ci95** | **continuous association spearman** | **p** |
| --- | --- | --- | --- | --- | --- | --- | --- |
| Complete task set | all included | full course | 88 | 0.783 | 0.69 to 0.88 | 0.47 | <0.001 |
| Single task only | only MOT | full course | 88 | 0.755 | 0.65 to 0.85 | 0.367 | <0.001 |
| Single task only | only RSVP | full course | 88 | 0.725 | 0.62 to 0.83 | 0.273 | 0.010 |
| Single task only | only Search | full course | 87 | 0.743 | 0.63 to 0.84 | 0.357 | <0.001 |
| Single task only | only Sweeps | full course | 79 | 0.707 | 0.58 to 0.82 | 0.317 | 0.004 |
| Single task only | only UFOV | full course | 74 | 0.675 | 0.54 to 0.80 | 0.268 | 0.021 |
| Leave-one-task-out | excluding MOT | full course | 88 | 0.78 | 0.68 to 0.88 | 0.468 | <0.001 |
| Leave-one-task-out | excluding RSVP | full course | 88 | 0.77 | 0.67 to 0.87 | 0.451 | <0.001 |
| Leave-one-task-out | excluding Search | full course | 88 | 0.754 | 0.66 to 0.85 | 0.435 | <0.001 |
| Leave-one-task-out | excluding Sweeps | full course | 88 | 0.749 | 0.65 to 0.85 | 0.406 | <0.001 |
| Leave-one-task-out | excluding UFOV | full course | 88 | 0.766 | 0.67 to 0.86 | 0.427 | <0.001 |
| Complete task set | all included | first 25% | 71 | 0.672 | 0.54 to 0.79 | 0.307 | 0.009 |
| Leave-one-task-out | excluding MOT | first 25% | 69 | 0.654 | 0.53 to 0.78 | 0.264 | 0.028 |
| Leave-one-task-out | excluding RSVP | first 25% | 70 | 0.617 | 0.47 to 0.75 | 0.248 | 0.038 |
| Leave-one-task-out | excluding Search | first 25% | 71 | 0.64 | 0.50 to 0.77 | 0.246 | 0.039 |
| Leave-one-task-out | excluding Sweeps | first 25% | 71 | 0.656 | 0.51 to 0.78 | 0.279 | 0.018 |
| Leave-one-task-out | excluding UFOV | first 25% | 71 | 0.68 | 0.55 to 0.80 | 0.303 | 0.010 |
| Complete task set | all included | first 50% | 82 | 0.761 | 0.65 to 0.86 | 0.439 | <0.001 |
| Leave-one-task-out | excluding MOT | first 50% | 82 | 0.732 | 0.62 to 0.84 | 0.394 | <0.001 |
| Leave-one-task-out | excluding RSVP | first 50% | 82 | 0.743 | 0.63 to 0.85 | 0.413 | <0.001 |
| Leave-one-task-out | excluding Search | first 50% | 82 | 0.761 | 0.65 to 0.86 | 0.426 | <0.001 |
| Leave-one-task-out | excluding Sweeps | first 50% | 82 | 0.739 | 0.62 to 0.85 | 0.37 | <0.001 |
| Leave-one-task-out | excluding UFOV | first 50% | 82 | 0.744 | 0.63 to 0.85 | 0.374 | <0.001 |

### Table S7B. Intervention heterogeneity

Table S7B. Prognostic FSS main effects and treatment-interaction tests. Coefficients are fully standardised (change in SD of baseline-adjusted post-intervention EF per 1 SD higher FSS) from ordinary-least-squares models with HC3 robust standard errors. Interaction tests are exploratory and underpowered. Main-effect p values marked [see note] were not stored by the source analysis for subgroup models.

| **cohort subgroup** | **comparison** | **n** | **prognostic FSS beta** | **ci95** | **main effect p** | **interaction term** | **interaction p** | **partial R2** | **robust se** |
| --- | --- | --- | --- | --- | --- | --- | --- | --- | --- |
| BREATHE, all participants | 3 randomised arms | 88 | 0.3 | 0.11-0.50 | 0.002 | FSS x arm (omnibus) | 0.38 | 0.023 | HC3 |
| BREATHE, MCI only | 3 randomised arms | 60 | 0.34 | 0.08-0.60 | [see note] | FSS x arm (omnibus) | 0.397 | 0.031 | HC3 |
| BREATHE, CCT+RFB arm (mixed status) | cognitively unimpaired vs MCI | 51 | 0.28 | 0.01-0.55 | [see note] | FSS x cognitive status | 0.619 | 0.003 | HC3 |
| PSOPT | CCT+interface vs mental-leisure control | 40 | 0.33 | -0.04-0.71 | 0.083 | FSS x arm | 0.46 |  | HC3 |

### Supplementary Figures

#### Figure S1. Participant flow and recording quality

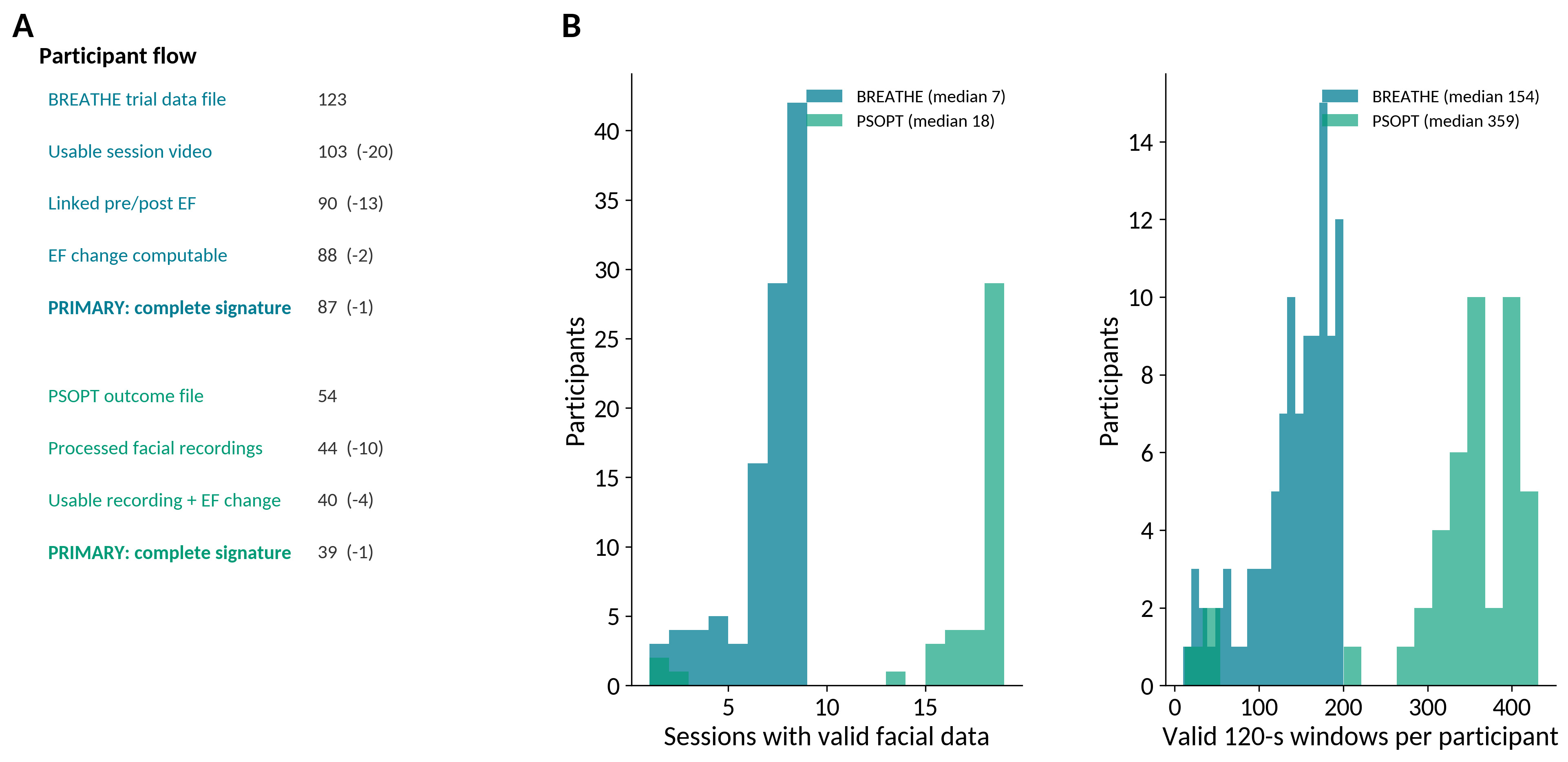

Figure S1. (A) Participant flow for the BREATHE development cohort and the PSOPT external-validation cohort, with exclusions at each stage. (B) Distribution of the number of sessions contributing valid facial data per participant. (C) Distribution of the number of valid 120-second windows per participant. A window was valid if at least one sampled frame yielded action-unit output. Recording quality metrics are those used in quality control.

#### Figure S2. Validation diagnostics

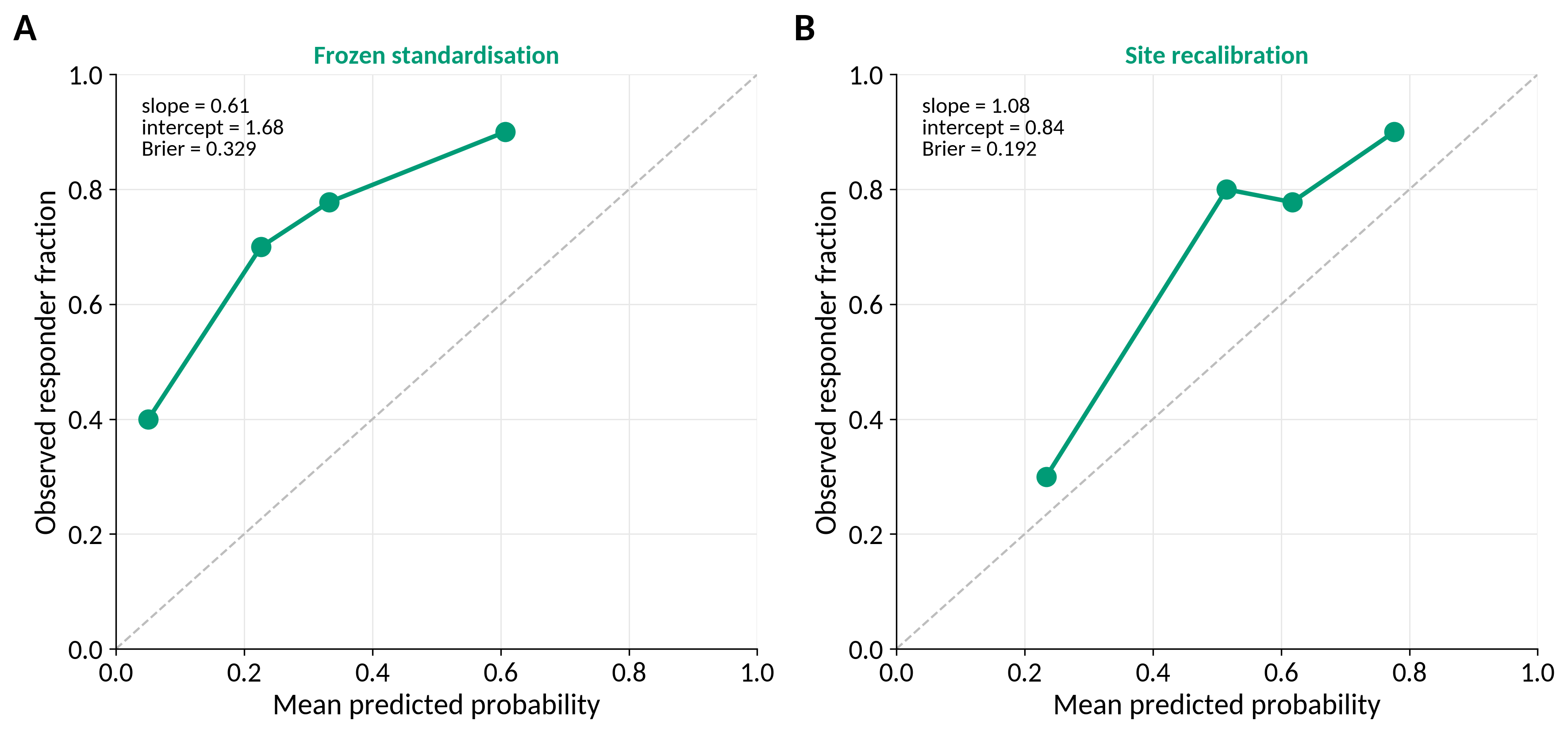

Figure S2. (A) BREATHE full-pipeline permutation distribution: the complete development pipeline, including feature screening, deduplication and ranking, was repeated for each of 1,000 participant-level outcome permutations. The observed statistic is the fixed-signature leave-one-subject-out AUROC (0.80), not the nested estimate. (B) PSOPT conditional permutation distribution: PSOPT outcome labels were permuted 1,000 times with all model parameters fixed. (C) PSOPT calibration under frozen development standardisation and (D) after outcome-independent site recalibration; points are quartiles of predicted probability, the dashed line is perfect calibration, and calibration slope, intercept and Brier score are shown. One-sided permutation p values were computed as (b + 1) / (B + 1).

#### Figure S3. Exploratory robustness

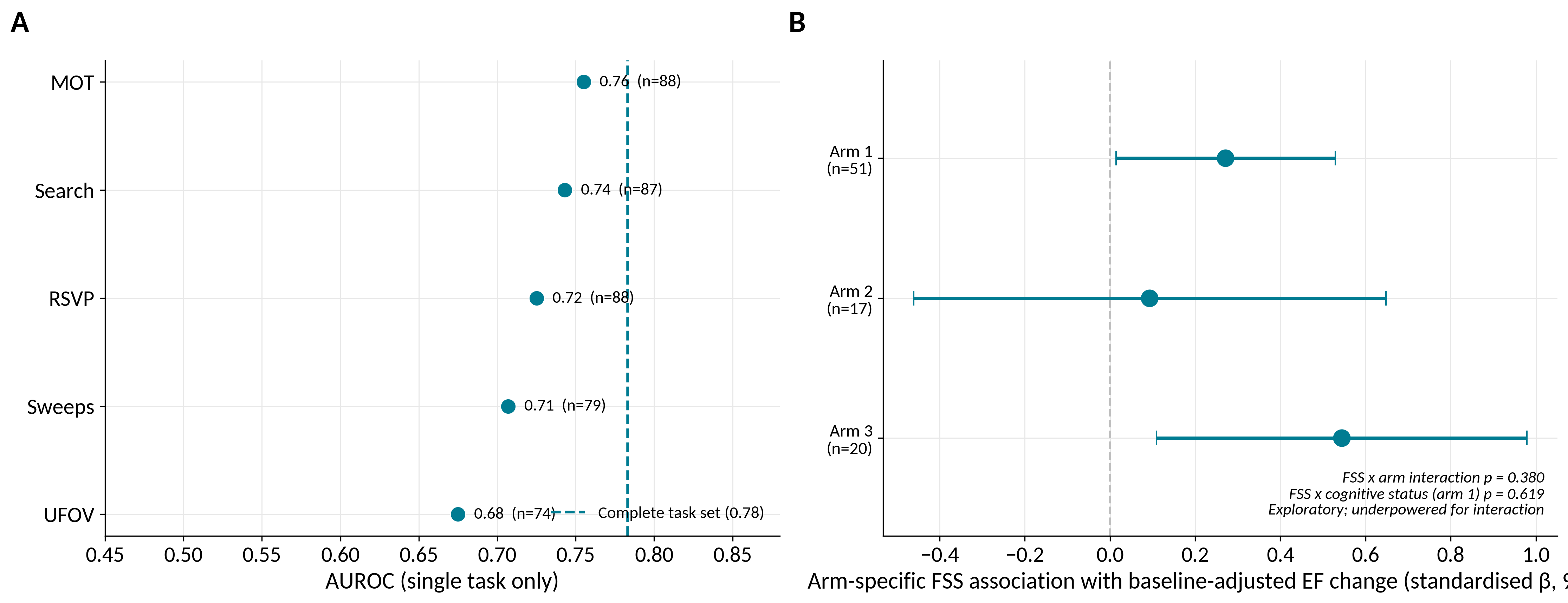

Figure S3. (A) Discrimination when the fixed three-feature signature was reconstructed from each BREATHE cognitive challenge alone, with the complete-task-set reference shown as a dashed line; sample sizes differ because features were not computable from every task for every participant. (B) Arm-specific associations between the FSS and baseline-adjusted EF change (fully standardised coefficients with 95% confidence intervals from HC3 robust standard errors), together with the omnibus FSS-by-arm and FSS-by-cognitive-status interaction tests. Both panels are exploratory; the interaction tests are underpowered and no arm-specific claim is made.
